# Diarylamidine activation of a brachiopod DEG/ENaC/ASIC channel

**DOI:** 10.1101/2024.08.26.609674

**Authors:** Josep Martí-Solans, Aina Børve, Andreas Hejnol, Timothy Lynagh

## Abstract

Diarylamidines are a group of widely used small molecule drugs. One common use of diarylamidines is their pharmacological inhibition of ligand-gated cation channels, including tetrameric ionotropic glutamate receptors and trimeric degenerin/epithelial sodium channel channel/acid-sensing ion channels (DEG/ENaC/ASICs). Here, we discover a DEG/ENaC/ASIC channel from the brachiopod (lamp shell) *Novocrania anomala*, at which diarylamidines act as agonists. The channel is closely related to bile acid-gated, pH-gated, and peptide-gated channels but is not activated by such stimuli. We describe activation of the channel by diminazene, DAPI, and pentamidine, examine several biophysical and pharmacological properties, and briefly explore the molecular determinants of channel activity with site-directed mutagenesis. We term this channel the diarylamidine-activated sodium channel (DiaaNaC).

## Introduction

Diarylamidines, such as pentamidine, diminazene, and 4′,6-diamidino-2-phenylindole (DAPI), are a class of chemical compounds distinguished by the presence of two aromatic amidine groups joined by a hydrophobic link. Diarylamidines are commonly utilized as antiparasitic drugs and cell nucleus dyes due to their capacity to strongly bind DNA AT-rich regions and interfere with DNA and RNA and thus protein synthesis (1). Interestingly, in addition to these capacities, diarylamidines are used as ion channels blockers. For example, various ionotropic glutamate receptors (iGluRs), which are mostly non-selective cation channels, are inhibited by diminazene, pentamidine, and/or DAPI(2-4) The inhibitory effect of diarylamidines is not restricted to iGluRs, as several members of the Degenerin/Epithelial Na^+^ Channel/Acid-Sensing Ion Channel (DEG/ENaC/ASIC) family of Na^+^ channels are also inhibited by diarylamidines (5-7). Indeed, diminazene is emerging as a widely used DEG/ENaC/ASIC blocker, because it may be more of a genuine pore blocker (i.e. plugs the narrowest part of the pore) of more numerous DEG/ENaC/ASIC channels than amiloride, which has traditionally been used to characterize channel pharmacology (5).

The DEG/ENaC/ASIC family includes a diverse range of ion channels that play crucial roles in various physiological processes, including mechanotransduction, ion homeostasis, and synaptic transmission (8). Members of the DEG/ENaC/ASIC ion channel family have a conserved trimeric structure, with three homologous subunits arranged three-fold symmetrically around a central pore (9). Each subunit has two transmembrane domains (TM1 and TM2) with a large extracellular region in between, and the N- and C-terminal domains are intracellular. TM1 and TM2 of each subunit form the channel pore, which is in most characterized cases moderately selective for Na^+^ over K^+^, usually with minimal divalent cation permeability(10). Several relatively well conserved amino acid residues in the mid-to upper part of the pore appear to determine the binding of cationic channel blockers, including diminazene, amiloride, and Ca^2+^ ions (5,7,11).

Computational docking and mutagenesis in rat ASIC1a suggest that both diminazene and amiloride plug the channel pore between rings of glycine residues (*TM2-G6’, TM2-G3’*) and aspartate residues (*TM2-D0’*)(5). An X-ray structure of chick ASIC1 instead shows amiloride molecules in two sites: one in the lateral portals, between adjacent subunits at the upper/lateral entrance to the pore and close to TM2-D0’, and another in a more distal extracellular site within single subunits(12). Cryo-electron microscopy structures of a more distantly related peptide-gated DEG/ENaC/ASIC channel in the presence of diminazene failed to resolve the drug, although a density was observed deep in the pore between TM2-G6’ residues, and mutagenesis of TM2 G6’ and to a lesser extent TM2-D0’ reduced potency of diminazene block(7). The upper half of TM1 and TM2 are intricately coupled to the extracellular domain, and are thus crucial for channel activation by extracellular stimuli, in addition to forming sites for drugs that modulate channel function (12-14).

While investigating the molecular basis by which various stimuli activate channels in a particular branch of the DEG/ENaC/ASIC family, we tested the activity of several channels from invertebrate bilaterian animals, including *Novocrania anomala*, a brachiopod (lamp shell). To our surprise, one channel was activated, not inhibited, by diarylamidines. Here, we describe the biophysical and pharmacological properties of this unusual DEG/ENaC/ASIC channel.

## Results

### Diarylamidines gate a DEG/ENaC/ASIC channel from the brachiopod *Novocrania anomala*

In a previous study (15), we performed a comprehensive phylogenetic analysis of the DEG/ENaC/ASIC family, identifying 11 DEG/ENaC/ASIC genes from the brachiopod (lamp shell) *N. anomala*. Notably, six of the *N. anomala* genes (asterisks in Fig. 1A) fall in the ASIC/BASIC/HyNaC branch within clade A DEG/ENaC/ASICs, one of two major clades that make up the family (15-18). In the present study, we expressed a *N. anomala* gene that is closely related to mammalian bile acid-sensing ion channels (BASICs) (Fig. 1A, red asterisk; Nano_50439 in (15)), in *Xenopus laevis* oocytes and measured its activity with two-electrode voltage clamp (TEVC). We sought to investigate if bile acids or protons, agonists of closely related BASIC and ASIC channels, would activate the *N. anomala* channel. Ursodeoxycholic acid (UDCA) and protons (pH 4), however, activated very little current—73.1 ± 10 nA and 64.6 ± 9 nA, respectively (Fig. 1B; n = 7). This suggests that, in contrast to closely related channels, the *N. anomala* channel does not significantly respond to these ligands.

**Fig. 1.**
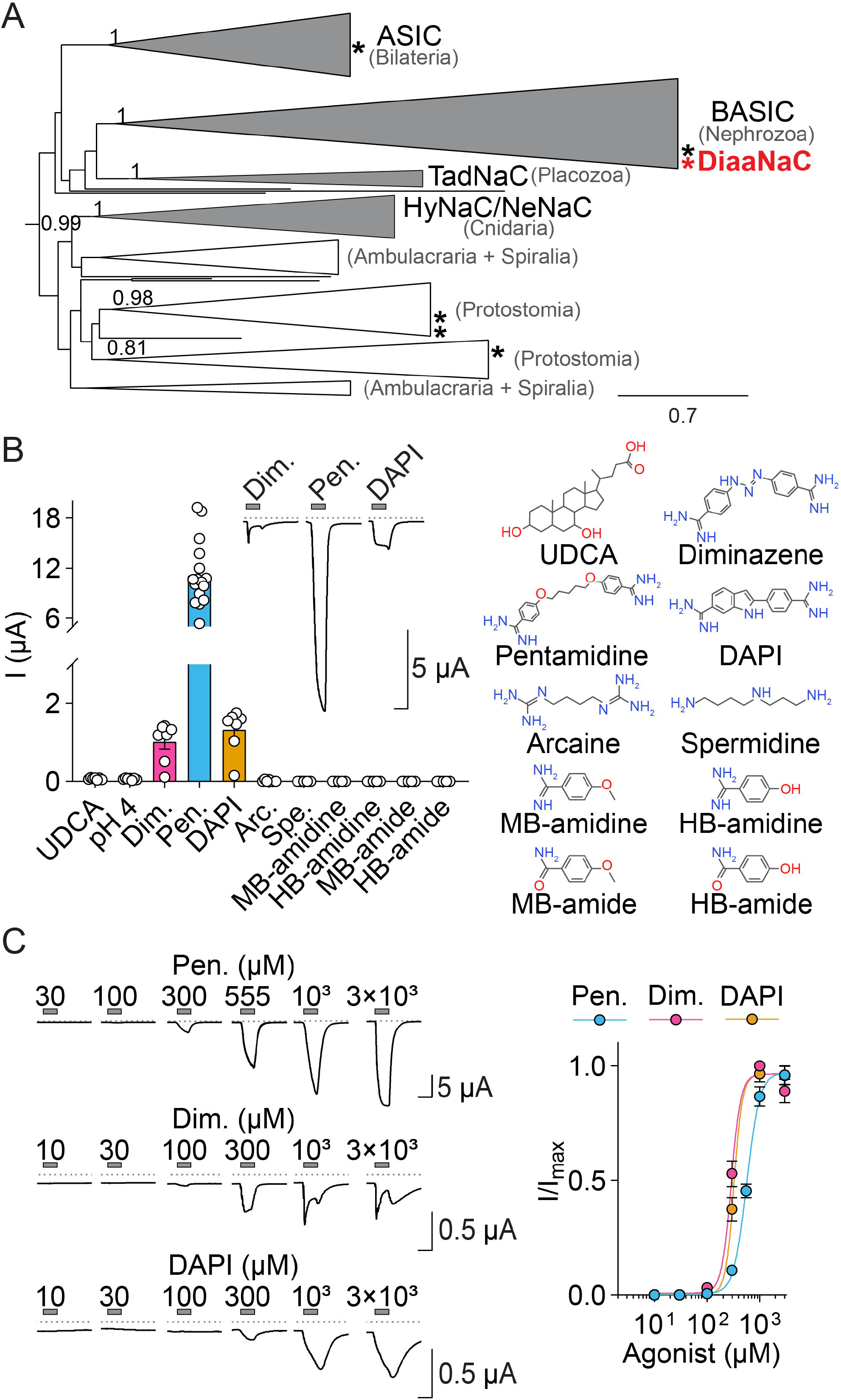
DiaaNaC channel is activated by diarylamidines. (A) ASIC/BASIC/HyNaC branch from the phylogenetic tree of the DEG/ENaC/ASIC family adapted from (15). Groups with functional data are shown in grey and groups without functional data in white. The taxonomic clades represented in each group are indicated in grey. Asterisks indicate *N. anomala* genes and red asterisk the gene characterized in this study (DiaaNaC). (B) *Left*, current activated by indicated compounds: UDCA, ursodeoxycholic acid; Dim., diminazene; Pen., pentamidine; DAPI, 2-(4-carbamimidoylphenyl)-1H-indole-6-carboximidamide; Arc., arcaine, Spe., spermidine; MB-amidine, 4-Methoxybenzamidine hydrochloride; HB-amidine, 4-Hydroxybenzamidine hydrochloride; MB-amide, 4-Methoxybenzamide; HB-amide, 4-Hydroxybenzamide. UDCA was tested at 2 mM and pentamidine at 300 μM, the other compounds at 1 mM. *Middle*, Example current traces of *Xenopus* oocytes expressing DiaaNaC. Grey dashed line indicates zero current level. *Right*, chemical structures (non-ionized form) of the compounds used in this study. (C) *Left*, Example current traces from DiaaNaC activated by different diarylamidine concentrations in absence of Ca^2+^. *Right*, mean (± SEM) normalized current amplitude in response to increasing diarylamidine concentrations for DiaaNaC in the absence of Ca^2+^.

Channels of the ASIC/BASIC/HyNaC branch show variable degrees of block by the antiparasitic agent diminazene (6,19,20). Considering this, we investigated the effects of diminazene on the *N. anomala* channel, which in the absence of Ca^2+^ in the extracellular solution showed a modest constitutive current of 343 ± 70 nA (n = 12). We attempted to inhibit this current by applying diminazene (1 mM). Unexpectedly, the application of diminazene activated substantial and reversible inward currents of 1.0 ± 0.2 μA (n = 7), without inhibiting the constitutive current (Fig. 1B). To investigate the specificity of this effect, we tested channel activation by other diarylamidines. Application of pentamidine (300 μM), which contains a flexible aliphatic chain, resulted in 12 times greater current amplitude than iminazene, whereas DAPI (1 mM) resulted in 1.3 times greater current amplitude than diminazene (Fig. 1B; n = 7–9). Therefore, we named this channel DiaaNaC (<u>Dia</u>rylamidine-activated <u>Na</u>^+^ <u>C</u>hannel).

We investigated requirements for agonist activity further, first by testing two aliphatic polyamine compounds, arcaine, which is a recognized ASIC3 modulator (21), and spermidine; however, neither activated DiaaNaC (Fig. 1B; n = 5-7), suggesting that the arylamidine moieties are important for agonist activity. We therefore tested the activity of monoarylamidines or monoarylamides, but none of these activated the channel (Fig. 1B; n = 5), indicating that the presence of two arylamidine groups is most probably necessary for channel activation.

We determined the potency of the different diarylamidines at DiaaNaC by applying the drugs in increasing concentrations. Diminazene and DAPI (EC_50_ 287.1 ± 16.5 and 361.6.1 ± 44.2 μM, respectively) showed higher potency than pentamidine (591.6 ± 30.7 μM), despite much greater efficacy of the latter (Fig. 1B, C). Because of the greater amplitude and more complete deactivation of pentamidine-gated currents, we used this drug in further studying the channel.

### DiaaNaC is inhibited by Amiloride and Ca^2+^

Diminazene and the diuretic drug amiloride, a cationic small molecule, compete for a similar binding site in the pore of ASIC and BASIC channels (5,6). Therefore, we investigated if amiloride could activate DiaaNaC like diarylamidines, but the application of amiloride (1 mM) activated no current in DiaaNaC-expressing oocytes (Fig. 2A). Instead, when co-applied with pentamidine, amiloride inhibited pentamidine-gated currents (IC_50_ of 1.1 ± 0.1 mM; n = 5). The presence of amiloride shifted the pentamidine concentration-response curve rightward (EC_50 + Ami._ = 3.6 ±0.9 mM; n = 5) but higher concentrations did not appear to relieve the inhibition by amiloride (Fig. 2B), indicating that the inhibition is not classically competitive (22), suggesting that amiloride inhibits via a different site than pentamidine. We questioned if amiloride inhibits via a site in the DiaaNaC channel pore and therefore tested inhibition at a depolarized membrane potential, which should weaken binding of such a positively charged molecule in the channel pore. However, the percentage of amiloride block did not change significantly when membrane voltage was raised from -100 mV to +10 mV (Fig. 2C), suggesting that amiloride block in DiaaNaC is not strongly voltage-dependent therefore not via a site that is deep in the electric field across the membrane.

**Fig. 2.**
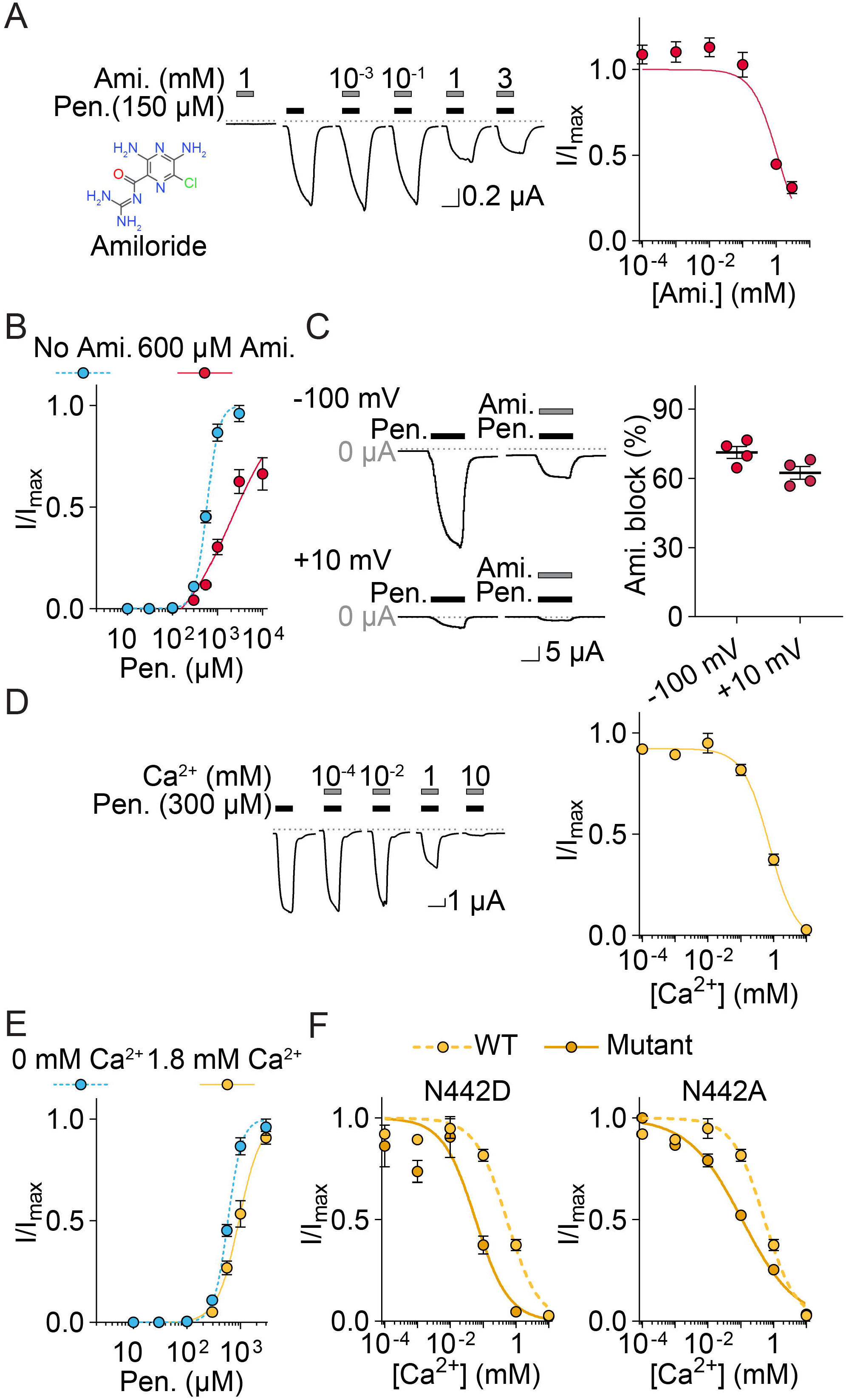
Amiloride and Ca^**2+**^ **block DiaaNaC**. (A) *Left-to-right*, amiloride structure (non-ionized form), example pentamidine (150 μM) -gated currents in increasing amiloride concentrations, and mean (± SEM, n=4) normalized current amplitude in increasing amiloride concentrations. (B) Mean (± SEM, n=4) normalized pentamidine-gated current amplitude alone or with 600 μM amiloride. (C) *Left*, example, and *right*, mean (± SEM, n=4) pentamidine (300 μM) -gated currents alone or with 1 mM amiloride at -100mV or +10mV membrane potential. (D) *Left*, example, and *right*, mean pentamidine (300 μM) -gated currents in increasing Ca^2+^ concentrations. (E) Mean (± SEM, n=4) normalized pentamidine-gated current amplitude without Ca^2+^ or with 1.8 mM Ca^2+^. (F) Mean (± SEM, n=4) normalized pentamidine (EC_10_ concnetration ) -gated current amplitude in response to increasing Ca^2+^ concentrations for indicated DiaaNaC mutants.

Next, we tested for potential inhibition of DiaaNaC by Ca^2+^, as inhibition by this divalent cation is broadly conserved in the ASIC/BASIC/HyNaC branch (20,23-25). After measuring the amplitude of pentamidine-gated currents of DiaaNaC at various Ca^2+^ concentrations, we found an apparent IC_50_ of 0.5 ± 0.1 mM (Fig.2B; n = 6) for Ca^2+^. This suggests that at physiological concentrations of extracellular Ca^2+^ (1.8 mM), DiaaNaC is largely inhibited. To investigate the antagonistic relationship of pentamidine and Ca^2+^, we measured the pentamidine concentration-response relationship at physiological Ca^2+^ concentrations and compared it to that in the absence of Ca^2+^. Ca^2+^ slightly shifted the concentration-response curve to the right (EC_50 + Ca2+_ = 986.3 ± 108.3; n = 6), and high concentrations of pentamidine could overcome the inhibitory effect of Ca^2+^, which we tentatively interpret as pentamidine and Ca^2+^ competing for the same binding site. The negatively charged TM2 D0’ residue in the upper half of the rat ASIC1a pore is crucial for diminazene and Ca^2+^ binding (5,11), and the homologous position in DiaaNaC is instead a neutral but isosteric asparagine residue, TM2 N0’ (N442). When we mutated TM2 N0’ to aspartate or to small, non-polar alanine, the DiaaNaC IC_50_ for Ca^2+^ shifted to the left in both cases (IC_50; N442D_ = 0.06 ± 0.02 mM and IC_50; N442A_ = 0.1 ± 0.02 mM). This is inconsistent with Ca^2+^, and by extrapolation pentamidine, binding to this site in the DiaaNaC pore.

### Channel properties of DiaaNaC

We next investigated if pentamidine itself permeates the DiaaNaC channel, which could contribute to the inward currents in high extracellular concentrations of pentamidine, a divalent cation at physiological pH due to its two amidine groups (26). If pentamidine permeates DiaaNaC, the membrane potential at which the current reverses direction, or reversal potential (E_rev_), should shift towards more positive potentials in increased extracellular pentamidine concentrations. We tested this by measuring current-voltage relationships at two different concentrations or pentamidine (150 and 450 μM) at membrane potentials from -80 mV to 100 mV (Fig. 3A). As we saw no significant shift in E_rev_ with increasing pentamidine concentration (E_rev,150μM_ = 53.5 ± 5.0 mV and E_rev, 450μM_ = 57.6 ± 3.4 mV; n = 5), despite large changes in current amplitude, we conclude that pentamidine does not permeate the channel.

**Fig. 3.**
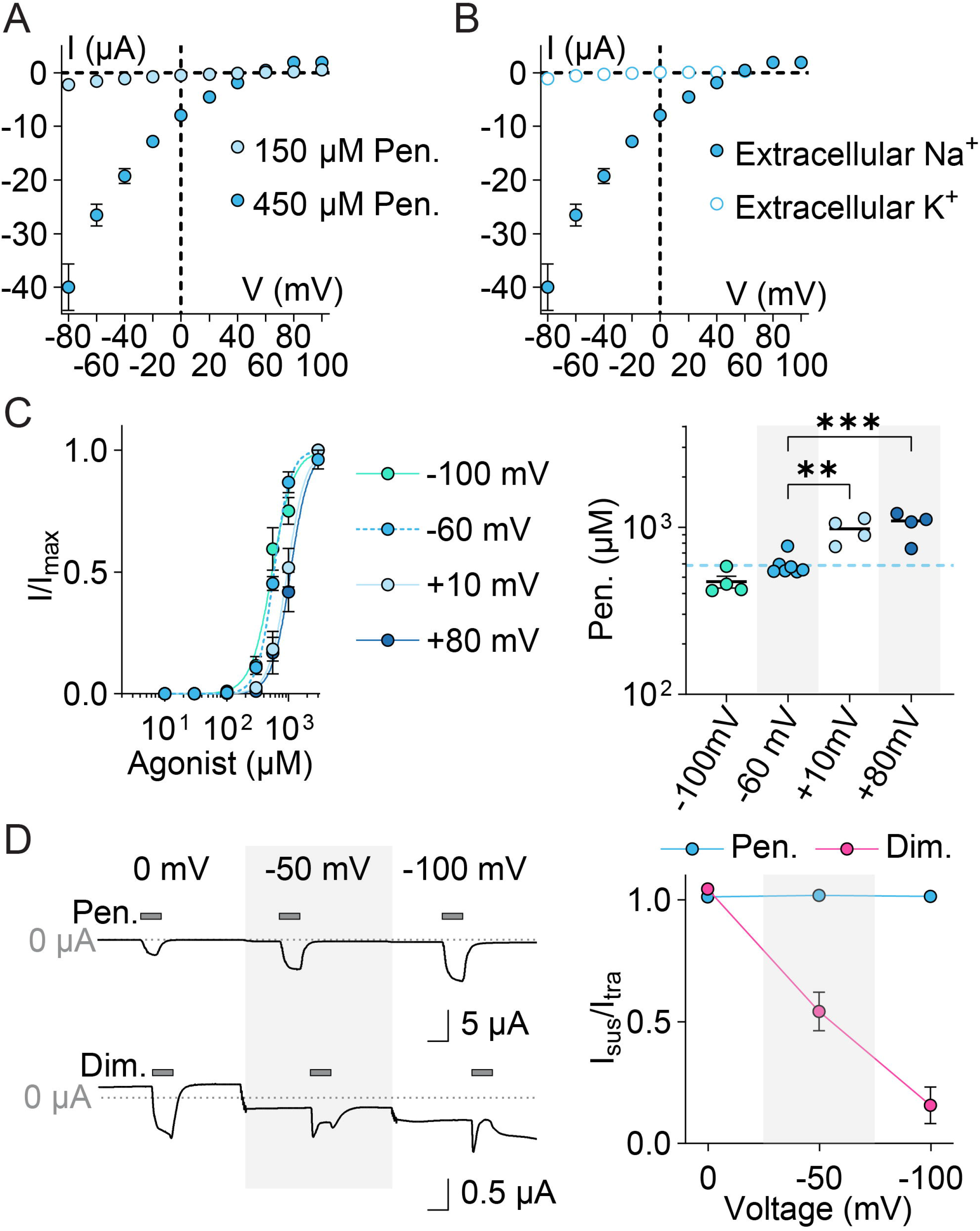
DiaaNaC channel properties. (A) Mean (± SEM, n=5) pentamidine (150 and 450 μM) - gated current at different membrane potentials (n=5). (B) Mean (± SEM, n=5) pentamidine-gated current at different membrane potentials in extracellular NaCl or KCl (n=5). (C) *Left*, mean (± SEM, n=4) normalized pentamidine-gated current amplitude in response to increasing concentrations of pentamidine at different membrane potentials (n = 4). *Right*, mean (± SEM, n=4-7) pentamidine EC_50_ compared by one-way ANOVA with Dunnett’s multiple comparisons test: significantly different than EC_50_ at -60mV marked with ** (p < 0.005) or *** (p < 0.0005). (D) *Left*, example 1 mM pentamidine (top) or diminazene (bottom) gated currents at indicated membrane potentials. *Right*, mean (± SEM, n=5) sustained current (I_sus_) amplitude normalized to transient current (I_trans_) amplitude at different membrane potentials. (E) *Left*, example, and *right*, mean (± SEM, n=4) normalized current amplitude in response to increasing diminazene concentrations at oocytes expressing indicated constructs.

We next considered permeation of monovalent cations through DiaaNaC. With typical high-Na^+^ extracellular solutions (in the absence of Ca^2+^), we measured a relatively positive E_rev_ of 57.6 ± 3.4 mV (n = 5), suggesting that the currents through this channel are predominantly carried by Na^+^. When K^+^ was the only monovalent cation in our extracellular solution, pentamidine-gated currents were tiny (Fig. 3B). This made it difficult to measure E_rev_ in this solution, but we estimated that E_rev_ = -12.2 ± 4.6 mV (n = 5) (Fig. 3B), a substantial shift in E_rev_. This indicates that DiaaNaC strongly prefers Na^+^ to K^+^, in terms of both permeability and conductance.

We next evaluated the effect of membrane potential on the apparent affinity of pentamidine for DiaaNaC. If the pentamidine binding site is in in the transmembrane region, as is the case for diarylamidines in other DEG/ENaC/ASIC channels (5,7), changes in membrane potential should affect the affinity of the positively charged drug for DiaaNaC. Therefore, we assessed apparent affinity via concentration-response experiments at four different voltages (Fig. 3C). When membrane potential was positive (i.e., +10 or +80 mV), the affinity of pentamidine for DiaaNaC decreased (EC_50, +10mV_ = 959.0 ± 80.7 and EC_50, +80mV_ = 1034 ± 101.2; n = 4) compared to reference (EC_50, -60mV_ = 591.6 ± 30.7; n = 7). Consistent with this, apparent affinity of pentamidine for DiaaNaC tended to increase at a more negative membrane potential (EC_50, -100mV_ = 471.0 ± 38.4). These results show that pentamidine activity is weakly voltage-dependent. This could derive from pentamidine binding just within the electric field, e.g. near the top of the membrane-spanning channel, or from membrane electric potential affecting the machinery of pentamidine-induced channel gating.

Unlike pentamidine and DAPI, diminazene-gated currents rapidly decreased in amplitude during application of the drug (Fig. 3D), especially at high concentrations (Fig. 1B,C). We hypothesized that this may be from diminazene activating the channel via one site and blocking the channel pore via another. Consistent with this hypothesis, diminazene blocked close to 100% of the current at -100 mV, and block was progressively relieved with decreased negative potential (Fig. 3D). In contrast, we observed no such block by pentamidine, even at -100 mV.

Furthermore, we tested diminazene activation of mutant TM2 G6’S channels, in which a narrow part of the channel pore near the middle of the lipid bilayer is further obscured by the addition of hydroxyl side chains. In this mutant, no channel block was observed, current amplitude was much greater than at WT channels, and the concentration-response relationship for activation was shifted to higher concentrations (Fig. 3E diminazene EC_50_ = 1270 ± 10 μM, n=3, compared to 287 ± 17 at WT). Together, these data point toward two facts about diarylamidine activity. Firstly, the agonist concentration-response relationship, as assayed by peak currents, underestimates the true agonist EC_50_ in the case of agonists that also block the channel. And secondly, diminazene and pentamidine share an agonist site that is either near the top or outside of the level of the membrane, and diminazene also binds to an inhibitory site deep in the pore.

### Molecular determinants of pentamidine activity

Finally, we sought to establish which amino acid residues determine activation of DiaaNaC by diarylamidines. We didn’t test all possible sites on the protein, but we tested potential roles of several residues in or near the top of the channel pore (Fig. 4A), as pentamidine potency was weakly voltage-dependent, and homologous residues in other DEG/ENaC/ASIC channels interact with diminazene and/or Ca^2+^and/or are important for channel gating, ion conduction, or modulation by drugs (5,11,12,14,27). We tested pentamidine activity at 11 single-mutant DiaaNaC channels carrying much larger or smaller side chains in this region, hypothesizing that mutations altering the size of the pore or the lateral portal would affect pentamidine potency if it acts via this region.

**Fig. 4.**
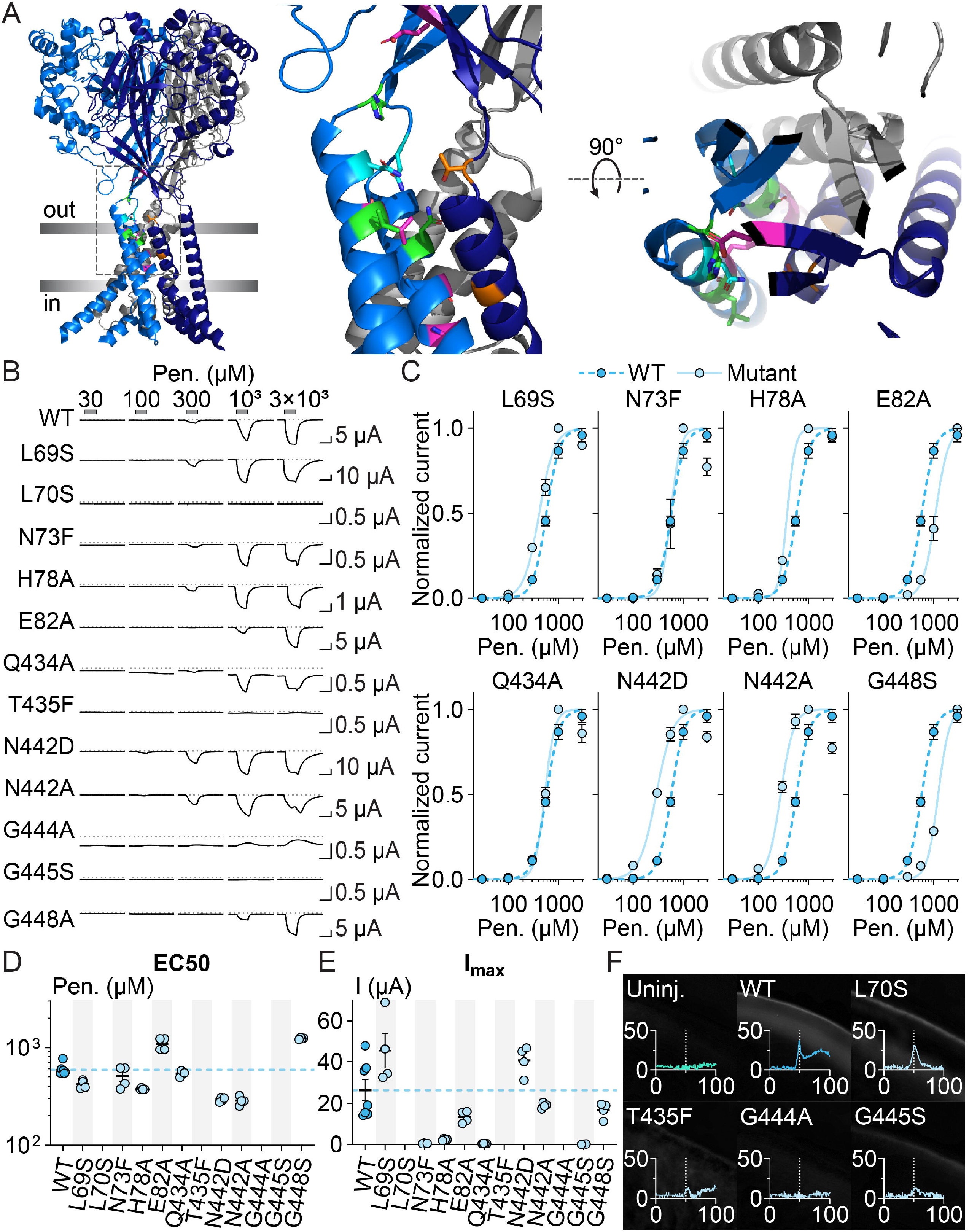
Potential determinants of pentamidine activity. (A) Side view (left and middle) and top-down view (right) of DiaaNaC model. Selected residues are indicated, with primer numbering for TM2. (B) Example current responses to pentamidine at oocytes expressing indicated DiaaNaC constructs. X-axis scale bars: 5 s. (C) Mean (± SEM, n=4) normalized pentamidinegated current amplitude for DiaaNaC mutants. (D) Mean (± SEM, n=4) pentamidine EC_50_ for indicated mutants. (E) Mean (± SEM, n=4) maximal pentamidine-gated current amplitude. (F) Immunolabeling of c-Myc tag in DiaaNaC mutants. Micropgraphs shown, and y, grey value, is plotted against x, distance from out-to-in (μM). White dashed line indicates presumptive cell membrane.

Three mutations, TM2 G2’A, TM2 G3’S, and TM2 T-7’F, abolished cell surface expression, as assayed by immunolabeling a C-terminal myc tag in our DiaaNaC construct (similar results on three oocytes each), in addition to the absence of pentamidine-gated currents (n = 5, Fig. 4B-F). Four mutations along the “left-hand side” of the portal had little or no effect on pentamidine potency, including TM1 N73F and TM2 Q-6A (no change in EC_50_), and TM1 L69S and post-TM1 H78A (<twofold decrease in EC_50_) (Fig. 4E). Three mutations caused moderate to extreme decreases in pentamidine potency. These included post-TM1 E82A and TM2 G6’S (∼twofold increase in EC_50_) and TM1 L70S, which abolished responses even to 3 mM pentamidine, despite apparent cell surface expression that was similar to WT (Fig. 4D-F). The two mutations that earlier increased the inhibitory potency of Ca^2+^ also increased the agonist potency of pentamidine. TM2 N0’D and N0’A mutants showed EC_50_ values of 293.5 ± 7.1 and 285.7 ± 14.2 (both n=4), twofold decreases relative to WT (Fig. 4B-D). This indicates that pentamidine activation of DiaaNaC does not rely on the N442 side chain, although the side chain may be important for pentamidine-induced gating. Thus, the lateral portal and upper half of the channel pore include certain residues important for activation by pentamidine, but the most important residues for high agonist potency are deep in the pore (TM2 G6’) and on the side (TM1 L70) or top (post-TM1 E82) of the portal, making it difficult to conclude from these experiments if pentamidine or other diarylamidines bind here to activate the channel.

## Discussion

Diarylamidines block several ligand-gated cation channels, as exemplified by diminazene block of numerous channels of the ASIC/BASIC/HyNaC branch of the DEG/ENaC/ASIC family. Here we identify DiaaNaC, a channel from this branch that is activated by diarylamidines.

DiaaNaC retains block by diminazene, which is strongly voltage-dependent and is abolished by the TM2-G6’S substitution, both reminiscent of diminazene block of ASICs and more distantly related FMRFamide-gated Na^+^ channels (5,7). The potency of diminazene block in DiaaNaC was difficult to assess due to concomitant activation and block, but block seems to develop at high micromolar and low millimolar concentrations. This is less potent than at the other channels mentioned and may derive from the absence of carboxylate side chains at the TM2-D0’ position, which provide favorable interactions for the cationic blocker in other DEG/ENaC/ASIC channels (5,7). We didn’t establish why pentamidine and DAPI exert no channel block, but perhaps the more numerous polar atoms in the “middle” of the diminazene molecule make it more suited the channel pore.

The agonist effects of the compounds appear to be via a different, more external site. Although the TM2 G6’S substitution decreased agonist potency of pentamidine twofold, we doubt that this constricted part of the pore is the binding site for the agonist. Loosely consistent with this, we found that in activating large currents, pentamidine does not appear to permeate the channel, and its agonist activity is only weakly dependent on membrane potential. Furthermore, we also observed decreases in agonist activity upon the post-TM1 E82A mutation, much higher in the protein, above the lateral portals. The largest decrease in potency among mutants tested was the TM1 L70S mutation, at a position which likely contacts the surrounding lipid bilayer. One possibility is that these compounds, with predicted water/octanol partition coefficients of around two (28), open the channel by associated with both the lipid bilayer and the upper half of the channel and/or the lower part of the extracellular domain. Certainly, modulators of ASICs seem to act at least partially via this region, as implicated by mutagenesis results investigating allosteric inhibition by ibuprofen (14) and structural data capturing both amiloride in the lateral portal and the lowest tip of the large, non-canonical toxin agonist MitTx in this vicinity (12).

Other possible binding sites could be more distal parts of the extracellular domain, although these would mean that the weak voltage dependence of pentamidine activation is due to an effect on the gating or conducting machinery. Potential extracellular binding sites include the space between the peripheral “thumb domain” and the more external “finger domain” domain of each subunit, equivalent to the “acidic pocket” in ASICs, which seems to undergo substantial conformational changes in DEG/ENaC/ASIC activation and via which several peptide drugs modulate ASIC activity (29). One study reporting computational docking of diarylamidines to an ASIC structure suggested a site in the peripheral thumb domain, although this lacks experimental verification (19). Establishing the precise mechanism of agonist activity by these typical DEG/ENaC/ASIC inhibitors will require more study in future. It’s curious that agonist activity seems competitive with Ca^2+^ ions, which themselves are thought to act on both the channel pore and extracellular sites to modulate DEG/ENaC/ASIC activity (11,30).

The biological role of DiaaNaC in *N. anomala* is even more mysterious. We are unaware of naturally occurring diarylamidines, and none of the naturally occurring compounds we tested activated the channel. We therefore presume there is a native agonist that remains to be identified. Curiously, we see no orthologue of DiaaNaC in transcriptomes of other brachiopods we examined, including *Terebretalia transversa* and *Lingula anatina*. If agonist activity of diarylamidines is indeed unique for *N. anomala* DiaaNaC, then diminazene remains a good tool for inhibiting DEG/ENaC/ASICs in most native preparations.

## Experimental procedures

### Identification of *N. anomala* DiaaNaC, molecular biology, and chemicals

Transcript Nano_50439, renamed DiaaNaC here, was a *Novocrania anomala* transcript identified as a DEG/ENaC/ASIC gene in a recent phylogenetic analysis of ours (15). The DiaaNaC full-length coding sequence was amplified by PCR from *N. anomala* cDNA and sub-cloned via SalI and BamHI sites into a pSP64 (polyA) vector (P1241, Promega) modified to contain the *Xenopus laevis* globin 5’ and 3’ UTRs and a C-terminal Myc-tag linked to the final DiaaNaC residue via Gly-Ser linker (the BamHI site) (see (15) for full sequence). The cDNA sequence was deposited in NCBI GenBank accession number PP942623. Recombinant PCR with Phusion High-Fidelity DNA polymerase (F-549L, Thermo Fisher Scientific) was used to perform site-directed mutagenesis, as described previously. Sanger sequencing of the whole insert (Genewiz) corroborated the sequences of wild-type (WT) and mutant constructs. EcoRI (FD0274, Thermo Fisher Scientific) was used to linearize the cDNAs via a site shortly after the poly(A) sequence, and the mMESSAGE mMACHINE SP6 Transcription Kit (AM1340, Thermo Fisher Scientific) was used to synthetize the cRNA.

Diminazene aceturate (D7770), pentamidine isethionate (P0547), spermidine trihydrochloride (S2501) and standard chemicals were purchased from Merck. DAPI (D1306) and other standard chemicals were purchased from Fisher Scientific. Other chemicals were purchased from other companies: 4-Methoxybenzamidine hydrochloride (F300402), 4-Hydroxybenzamidine hydrochloride (F023564), 4-Hydroxybenzamide (F002826), 4-Methoxybenzamide (F240987) from Fluorochem; arcaine sulfate (A-285) from Alomone Labs; sodium ursodeoxycholic acid (sc-222407) from Santa Cruz Biotechnology. 2D chemical structures were drawn with PubChem Sketcher V2.4.

### Heterologous expression and detection of surface expression

12 ng of WT cRNA for most experiments with WT channels, and 48 ng of WT (for additional diminazene experiments and for comparing pentamidine potency with mutants) and mutant cRNA was injected into stage V/VI *Xenopus laevis* oocytes purchased from Ecocyte Bioscience (Dortmund, Germany). Before electrophysiological recordings or surface expression determination, the oocytes were cultured for three days at 18°C in 50% Leibowitz medium (L1518, Merck), supplemented with 0.25 mg/ml gentamicin, 1 mM L-glutamine, and 15 mM HEPES (pH 7.6). For surface expression detection, mouse anti-Myc tag monoclonal IgG1 antibody (MA121316, Fisher Scientific) was used to detect the myc tag attached to the 3’-end of all the constructs used in this work, and the protocol previously used in our lab was followed (31) using a Zeiss Axio Scope A1 microscope. The intensity of the expression signal was quantified by tracing a perpendicular line across the oocyte membrane and measuring the grey values with ImageJ software (32).

### Electrophysiological recordings and data analysis

Whole cell currents were recorded from oocytes by two-electrode voltage clamp using an OC-725D amplifier (Warner Instruments) controlled via an Axon Digitata 1550B interface and pClamp v11 (Molecular Devices), acquired at 1 kHz and filtered at 200 Hz. Currents were analyzed in pClamp v10.7 software (Molecular Devices) and additionally filtered at 10 Hz for display in figures. Oocytes were clamped at -60 mV, unless otherwise indicated, and continuously perfused with a bath solution containing (in mM): 140 NaCl, 1.8 BaCl_2_, and 10 HEPES. pH was adjusted to 7.5 with NaOH, HCl, or KOH, as appropriate. In most experiments, activating solution was applied to oocytes in between resting periods of at least 30-60 s. After retrieving current amplitude from pClamp, all data were analysed in Prism v9 (GraphPad Software). For current-voltage relationships where K^+^ replaced Na^+^, KCl replaced NaCl in the above bath solution.

### Structural Model

The DiaaNaC trimeric structure was predicted using AlphaFold3 on the Google Colab site (33). The model was visualized in Fig. 4 with PyMol v2.4 (Schrödinger).

## Other notes

### Author contributions

JMS: conceptualization, investigation, analysis, visualization, manuscript writing and editing. AB: investigation, manuscript editing. AH: contributed resources, manuscript editing. TL: conceptualization, analysis, manuscript writing and editing.

### Funding and additional information

This work was supported by The Research Council of Norway, project number 234817.

